# STOmics-GenX: CRISPR based approach to improve cell identity specific gene detection from spatially resolved transcriptomics

**DOI:** 10.1101/2022.12.08.519589

**Authors:** Jennifer Currenti, Liang Qiao, Rhea Pai, Saurabh Gupta, Costerwell Khyriem, Kellie Wise, Xiaohuan Sun, Jon Armstrong, Jordan Crane, Smita Pathak, Bicheng Yang, Jacob George, Jasmine Plummer, Luciano Martelotto, Ankur Sharma

**Author notes:** Correspondence (L.M.); (A.S.).

## Abstract

The spatial organisation of cells defines the biological functions of tissue ecosystems from development to disease. Recently, an array of technologies have been developed to query gene expression in a spatial context. These include techniques such as employing barcoded oligonucleotides, single-molecule fluorescence in situ hybridization (smFISH), and DNA nanoball (DNB)-patterned arrays. However, resolution and efficiency vary across platforms and technologies. To obtain spatially relevant biological information from spatially resolved transcriptomics, we combined the Stereo-seq workflow with CRISPRclean technology to develop the STOmics-GenX pipeline. STOmics-GenX not only allowed us to reduce genomic, mitochondrial, and ribosomal reads, but also lead to a ∼2.1-fold increase in the number of detected genes when compared to conventional Stereo-seq (STOmics). Additionally, the STOmics-GenX pipeline resulted in an improved detection of cell type specific genes, thereby improving cellular annotations. Most importantly, STOmics-GenX allowed for enhanced detection of clinically relevant biomarkers such as Alpha-fetoprotein (AFP), enabling the identification of two spatially distinct subsets of hepatocytes in hepatocellular carcinoma tissue. Thereby, combining CRISPRclean technology with STOmics not only allowed improved gene detection but also paved the way for spatial precision oncology by improved detection of clinically relevant biomarkers.

## INTRODUCTION

Tissues and cells display remarkable heterogeneity (1), as highlighted by single cell and bulk technologies (2, 3). In the last decade, single-cell RNA sequencing (scRNAseq) technology has provided unprecedented insights into cellular heterogeneity (4). However, these dissociation-based technologies lack the cellular context of tissue and therefore, new spatial technologies have emerged to address this limitation (reviewed in (5)) (6-10). Spatial technologies map gene expression data to tissue architecture, allowing for biological insights while maintaining cellular context (11, 12). Such technologies are rapidly evolving with one of the most recent participants combining *in situ* RNA capture with DNA nanoball-patterned arrays, creating spatially enhanced resolution omics-sequencing (Stereo-seq) by BGI. Stereo-seq technology enables large field-of-view spatial transcriptomics, capturing the whole transcriptome at a nanoscale resolution with spots approximately 220nm in diameter with a centre-to-centre distance of 500nm (11). This technology uses STOmics gene expression chips containing the Stereo-seq capture units. STOmics has led to several spatiotemporal transcriptomic atlases with the technology permitting the discovery of several novel biological insights in developmental and disease contexts (11, 13-15). These insights have been possible in part due to the single-cell resolution, high sensitivity, and large field of view of STOmics (11).

Whole transcriptomic profiles from scRNA-seq and certain spatial methods contain not only messenger RNA (mRNA) information but also consists of reads corresponding to ribosomal and mitochondrial RNA (16). Furthermore, the detection efficiency of scRNAseq techniques lies between 3-25% (17, 18) and is likely similar for spatial technologies utilizing MRna capture techniques. For many downstream analyses such as cell type identification and cell-cell interaction analysis, only MRna information is used while reads corresponding to ribosomal and mitochondrial RNA are filtered out during quality control steps. This is exemplified by a recent study that was able to elucidate neutrophil heterogeneity in the liver tumor immune microenvironments by filtering out 1,514 genes associated with mitochondria, heat-shock protein, and ribosome prior to sub-clustering (19). While probe-based methods (7, 20) could improve the detection of MRna from scRNA-seq and spatial methods, it adds additional customisation costs to experiments (5). Therefore, we hypothesize that a CRISPR based approach of depleting uninformative reads (corresponding to housekeeping genes and genomic reads) could improve the detection of biologically relevant genes. To this end, we applied Jumpcode’s CRISPRclean technology to the STOmics workflow to improve biologically relevant signals from this new spatial transcriptomics assay.

Combining STOmics and CRISPRclean technologies aims to reduce noise in resultant data, increase the number of genes (detection efficiency) and their associated read count, reducing the cost of performing this spatial technology, widening the accessibility and capability of this technology. To this end, we utilized the newly developed spatial technology STOmics with CRISPRclean (henceforth STOmics-GenX) and without (conventional STOmics; **Fig. 1A**). STOmics-GenX captures the whole transcriptome *in situ* at a nanoscale resolution while removing “noise”. CRISPRclean leverages Cas9 depletion with specifically designed guides to remove genes and transcripts that would otherwise be removed using *post-hoc* computational steps prior to data analysis. STOmics-GenX has an enhanced ability to detect genes at a higher coverage, broadening the potential for biological discoveries in a spatial context.

**Fig 1.**
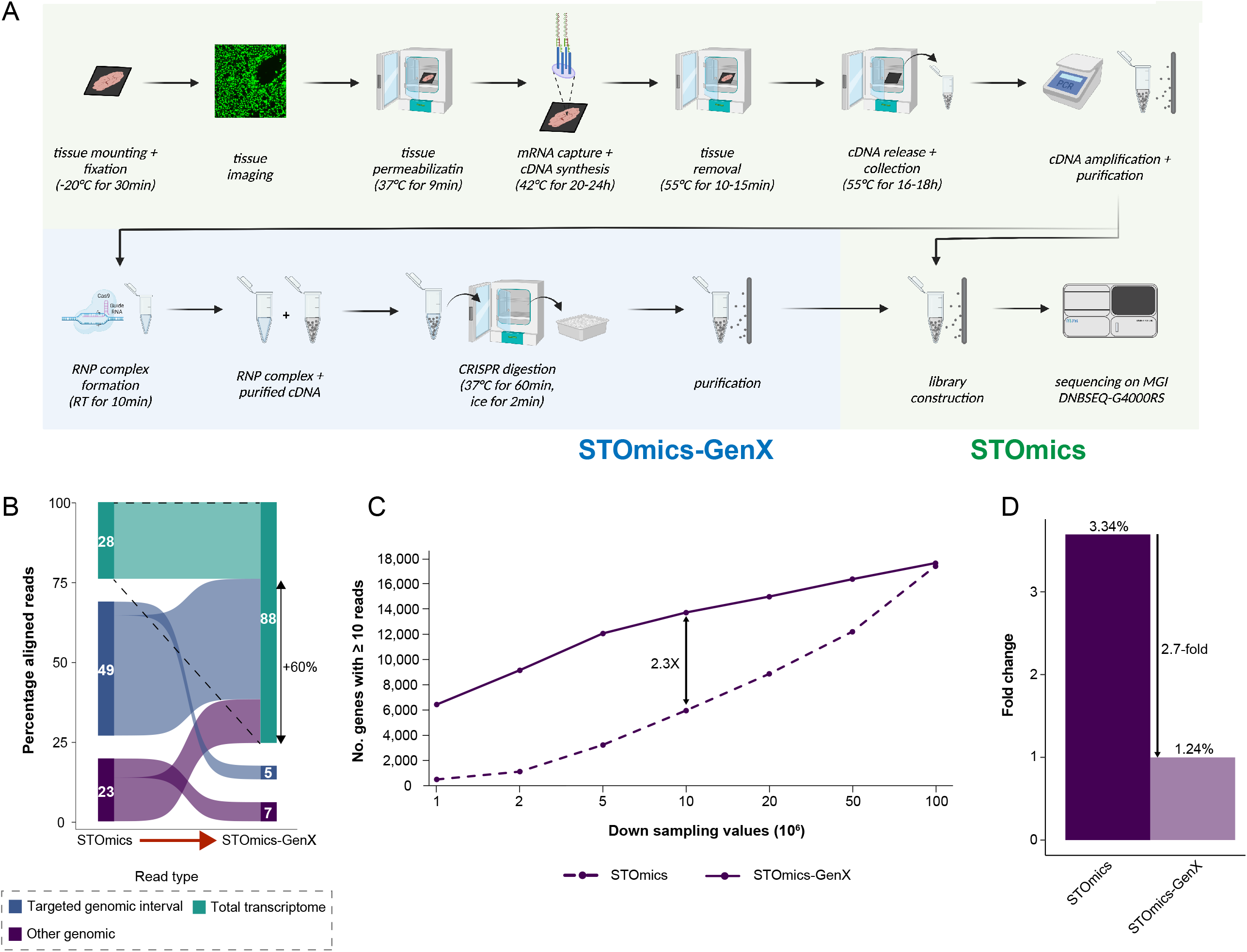
STOmics-GenX increases gene detection. **(A)** Purified cDNA from the conventional STOmics pipeline is utilized for CRISPRclean (STOmics-GenX) prior to library construction and sequencing. **(B)** Aligned reads with a valid co-ordinate identity showed a decrease in genomic reads and a subsequent increase in transcriptomic reads following STOmics-GenX. **(C)** Down sampling of aligned reads revealed an increase in detected genes with at least 10 reads using STOmics-GenX. **(D)** STOmics-GenX resulted in a 2.7-fold decrease in ribosomal and mitochondrial reads. The y-axis denotes the fold change relative to the depleted sample with ribosomal and mitochondrial read percentages indicated above respective bars.

## RESULTS

### STOmics-GenX increases conventional STOmics detection efficiency

A known trade-off with spatial methods exists between the number of genes and the capture efficiency, i.e. the proportion of reads aligning to transcripts of interest (5). While some spatial technologies use a probe-based method (7, 20) to improve the capture rate and thereby improve the read coverage for genes of interest, other methods such as STOmics (11) utilize an unbiased approach. Here we investigated whether the removal of housekeeping genes (including mitochondrial and ribosomal RNA) improves the detection of cell-type specific genes in STOmics. For this, we employed Jumpcode Genomic’s CRISPRclean technology which removes 349 genes associated with housekeeping and non-variable genes and genomic reads. Application of CRISPRclean with STOmics (STOmics-GenX) led to a reduction in targeted genomic reads from 49% to 5% (**Fig. 1B**).

To confirm if the re-distribution of aligned reads to the transcriptome translated to an increase in genes and in turn detection efficiency, we assessed library complexity between STOmics and STOmics-GenX. Reads with a valid co-ordinate identity (CID) were down sampled and the number of genes present with 10 or more reads was determined. Subsequently, an increase in the number of genes with 10 or more reads was observed using STOmics-GenX (**Fig. 1C**). Interestingly, at one million reads, STOmics-GenX detected 6,435 genes while STOmics detected 500 genes. Furthermore, for STOmics to detect a rough equivalent of 5,963 genes 10 million reads were required, at which point STOmics-GenX detected 13,730, a 2.3X increase. STOmics-GenX consistently revealed greater gene detection up until sequencing saturation at one billion reads, suggestive of an overall increase in detection efficiency. However, CRISPRclean is not only designed to reduce genomic reads, but also reduce the proportion of reads aligning to mitochondrial and ribosomal genes. Overall, a 2.7-fold decrease in aligned reads to ribosomal and mitochondrial genes was identified (**Fig. 1D**). Paired with increased detection efficiency, a greater capacity to interrogate biological questions likely exists with STOmics-GenX.

### STOmics methodology influences highly variable genes

Highly variable gene (HVG) analysis is predominantly utilized to comprehend biologically relevant genes and for cell-type annotations in scRNA-seq and spatial transcriptomics datasets (21). Therefore, we investigated the impact of reduction in genomic, ribosomal, and mitochondrial reads (as observed with STOmics-GenX) on the identification of HVGs and potential implications on cell-type annotations. HVGs were determined as having a residual variance above 1.4, resulting in 1,734 and 2,326 HVGs following STOmics and STOmics-GenX, respectively (**Fig. S1A**). HVGs only present in one methodology were identified, with 719 in STOmics (**Fig. S1B**) and 1,311 in STOmics-GenX (**Fig. S1C**) alone. Both methodologies **(Fig. S1D)** revealed 1,015 common HVGs, three of these being mitochondrial or ribosomal. Consistent with the reduction in mitochondrial or ribosomal reads was the identification of 21 mitochondrial or ribosomal genes as HVGs for STOmics alone (highlighted as red in **Fig. S1B**). Furthermore, these genes have higher expression in STOmics compared to STOmics-GenX, likely impacting their detection as HVGs.

### Improved detection of cell-type markers with STOmics-GenX

To determine if the reduction of mitochondrial or ribosomal genes overall and in HVGs for STOmics-GenX resulted in an improved ability to identify cell-type specific markers (epithelial, myeloid, endothelial, fibroblast, and lymphoid), we calculated the total expression of canonical cell subtype markers at bin50 (50 × 50 DNB, approximately 13 cells). Importantly, all major cell populations were identified by both methodologies at this resolution (**Fig. 2A**). However, when specifically investigating the expression of epithelial genes in these datasets from hepatocellular carcinoma tissue, we observed a marked increase in detection of epithelial genes in STOmics-GenX condition (**Fig. 2B-C**). Next, we investigated cell types associated with the tumor microenvironment, myeloid cells were detected in an additional 1,227 bins following STOmics-GenX (**Fig. 2D**) and while they displayed a similar count profile between both methodologies, we detected more bins with lower UMI counts in STOmics-GenX (**Fig. 2E**). Similarly, results were obtained for fibroblasts where an additional 732 bins were detected with STOmics-GenX (**Fig. 2F-G**). Endothelial cells were identified in 399 additional bins (**Fig. 2H-I**). Lastly, for lymphoid cells 199 more bins were identified with higher counts following STOmics-GenX (**Fig. 2J-K**). Consistently more bins containing canonical marker genes and with higher expression were identified following STOmics-GenX. These results suggest that depletion of housekeeping genes and genomic reads could improve the detection of genes with lower UMI counts in this spatial transcriptomics assay.

**Fig 2.**
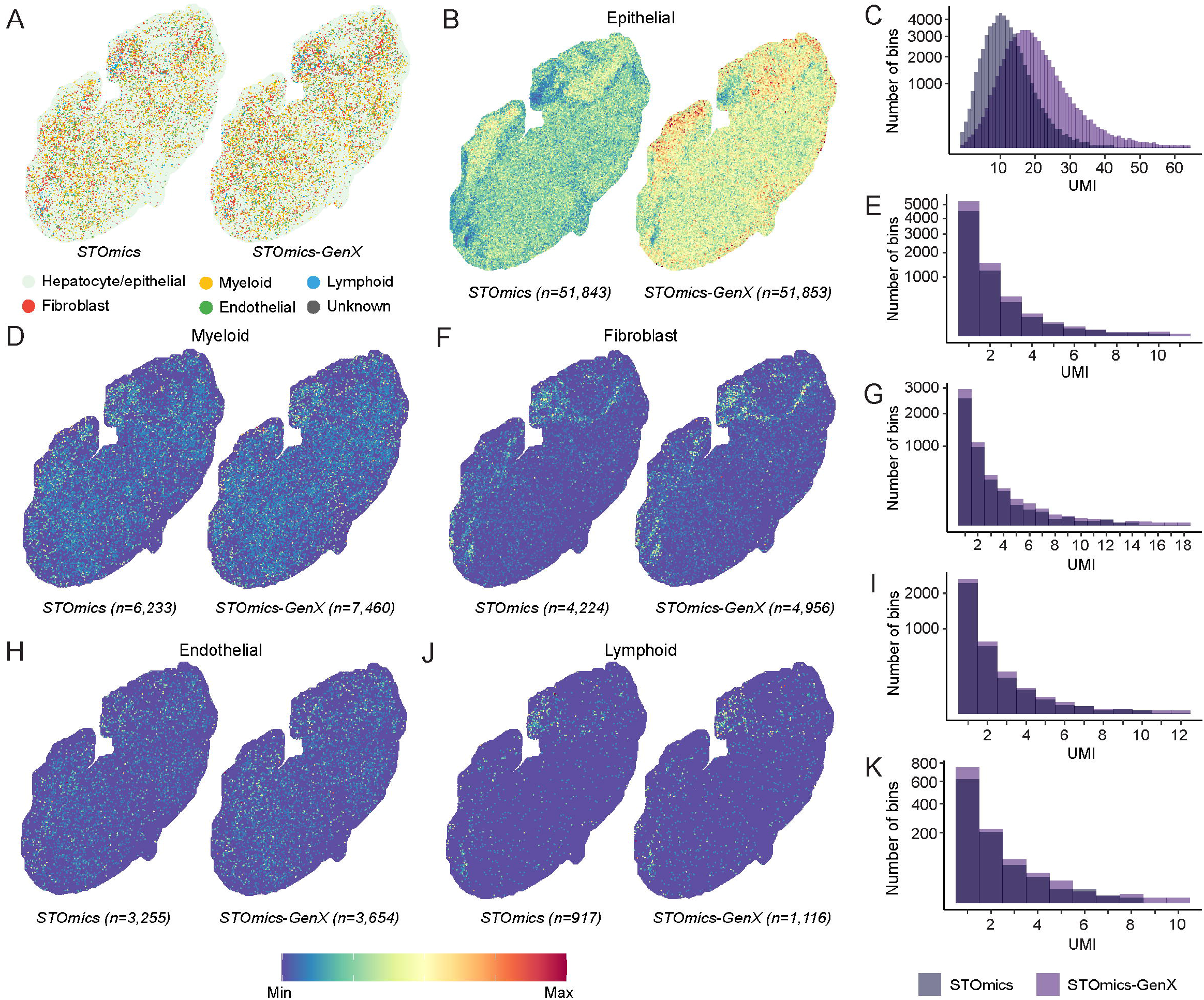
STOmics-GenX improves canonical gene expression. **(A)** The five major cell types can be annotated using both methodologies. **(B-C)** Hepatocytes/epithelial cells are identified by both methodologies; however, the expression of hepatocyte/epithelial cell markers is higher following STOmics-GenX. Total expression of canonical cell type markers was enhanced by STOmics-GenX for myeloid **(D-E)**, fibroblast **(F-G)**, endothelial **(H-I)**, and lymphoid **(J-K)** cells. Furthermore, these cells were consistently detected in more bins at higher UMI counts following STOmics-GenX compared to conventional STOmics.

### Implications for detecting clinical biomarkers

Since STOmics-GenX resulted in an improved detection of cell-type specific genes, including those with lower UMI counts, we next investigated whether STOmics-GenX also improved the ability to identify and detect clinically relevant biomarkers. Alpha fetoprotein (AFP) is one of the single most important biomarkers for the detection of hepatocellular carcinoma (HCC) in a clinical setting (22). Importantly, AFP levels could also predict early recurrence in HCC patients (23, 24). Similarly, haptoglobin (HP) is another key biomarker associated with overall survival in HCC patients (22, 25). In order to evaluate the potential implication of spatial transcriptomics in precision oncology, we investigated AFP (**Fig. 3A-B**) and HP (**Fig. 3C-D**) levels across both methodologies. Interestingly, we observed improved detection of AFP level by STOmics-GenX, suggesting potential benefits of CRISPRclean technology in improving the detection of clinically relevant biomarkers. Moreover, based on the expression of AFP and HP, we also identified two specific populations of hepatocytes (AFP^hi^HP^lo^ and AFP^lo^HP^hii^) with distinct spatial locations in tumor tissue. These results demonstrate the potential of STOmics-GenX in obtaining clinically relevant information from tumors while preserving their spatial context.

**Fig 3.**
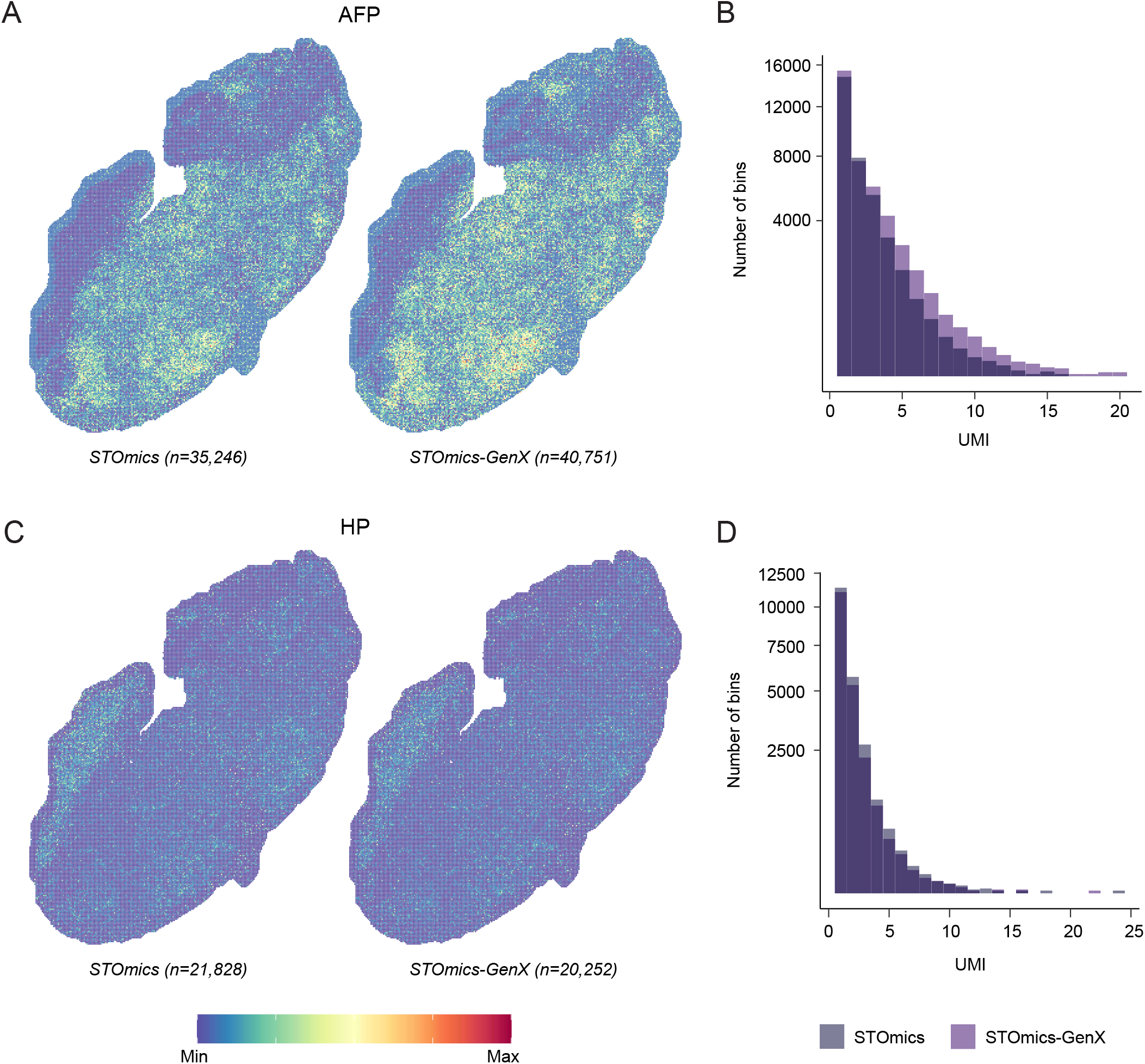
Spatial localization of cell populations. AFP expression was identified in an additional 5,505 bins following STOmics-GenX **(A)** and exhibited a slight shift towards higher overall expression **(B)**. HP expression was identified in a similar number of bins with similar expression between methodologies **(C-D)**.

## DISCUSSION

Single cell technologies have enabled the unbiased assessment of cellular heterogeneity, identity, and phenotypic properties at an unprecedented scale (26). Specifically, spatial transcriptomics allows for the interrogation of cellular heterogeneity in a spatial context, improving our understanding of cell-cell communications in health and disease (26). Most importantly, spatial transcriptomics allows for the identification of cellular ecosystems associated with development and disease progression (27). The analysis of both scRNAseq and spatial data involves several quality control steps, of which regression or complete removal of mitochondrial and/or ribosomal genes is common and is an inbuilt feature of the widely used pipelines Seurat (28) and Scanpy (29). The removal of these genes prior to clustering has enabled the detection of distinct cell subtypes (19) and may become a more common feature to identify sub-clusters from major cell types. As a large portion of scRNA-seq and spatial technologies centre around poly-A captured targets, detection of mitochondrial and ribosomal RNA is an integral part of the workflow (5). Recently, probe-based technologies have emerged which not only improve detection efficiency, but also circumvent the issues related to mitochondrial and ribosomal RNAs (7, 26). However, probe-based approaches require the customization of gene panels and therefore might not be ideal for larger panels due to cost barriers. An alternative approach could be to use a one-step method to remove housekeeping genes as well as mitochondrial and ribosomal RNAs, thereby increasing sequencing reads for biologically relevant genes. Recently, Jumpcode Genomics developed CRISPRclean, a technology utilizing Cas9 depletion with guides targeting genomic intervals alongside mitochondrial and ribosomal RNA to remove “noise” *in vitro*. This technology allows reads to be re-distributed to genes of interest and in turn, increasing detection efficiency.

Here, we investigated the utility of CRISPRclean technology in the conventional STOmics workflow (i.e. STOmics-GenX) and evaluated its impact on downstream analyses. STOmics-GenX led to a reduction in genomic reads, allowing reads to be re-distributed towards the transcriptome. As a result, more genes were detected at an equivalent and lower sequencing depth. In addition, we also observed a reduced presence of mitochondrial and ribosomal genes overall and as HVGs in STOmics-GenX. This translated to an improved ability to identify cell-type specific genes associated with immune and stromal cells in the tumor microenvironment. Therefore, STOmics-GenX allowed for increased gene detection while maintaining an unbiased approach. Aside from the increased efficiency, the most notable feature was the improved detection of clinically relevant biomarker such as AFP (22, 25, 30-33). Furthermore, this also led to the identification of two hepatocyte populations (AFP^hi^HP^lo^ and AFP^lo^HP^hi^) in HCC tissue with distinct spatial localization. Overall, these results indicate the power of an unbiased spatial analysis in discovering clinically relevant biological insights in the tumor microenvironment.

The increased efficiency of STOmics-GenX translates to a greater capacity to discover new biological insights with a marked reduction in sequencing cost. This reduction in cost permits a larger sample size, an important factor for identifying and validating clinical or biological biomarkers. Tumor heterogeneity is thought to be a driver of therapy failure and subsequent tumor recurrence (34). We have shown here that STOmics-GenX improves the ability to dissect tumor heterogeneity in HCC and this finding will likely extend to other cancer types and into the clinic. As such, STOmics-GenX is a more widely accessible spatial technology with the capability of detecting clinically relevant biomarkers, paving the way for precision oncology.

## MATERIALS AND METHODS

### Tissue preparation

Human hepatocellular carcinoma (HCC) samples were obtained from resected tissues and snap-frozen in Tissue-Tek O.C.T (Sakura, CA, USA) using pre-cooled isopentane. OCT-embedded tissues were stored long-term at -80°C. Cryosections were taken at a thickness of 8μm using a Leica GM3050 S cryostat.

### STOmics chips

STOmics chips are built utilizing Stereo-seq technology, DNA nanoball (DNB) patterned array technology with spatially barcoded probes, enabling the capture of the whole transcriptome *in situ* at a nanoscale resolution (https://www.stomics.tech/TranscriptomicsSet/). Resultant spatially resolved 3’ MRna libraries enable gene expression mapping of a tissue section.

### STOmics

STOmics was performed as described previously (11). Briefly, cryosections were mounted on the STOmics gene expression chip and fixed in pre-cooled methanol for 30min at -20°C. After fixation, tissues were stained with qubit ssDNA reagent (Invitrogen) and imaged on a Nikon Ni-E microscope utilising the FITC channel and 10x objective. Images were stitched together with 10% overlap to capture the entire stereo-seq chip. Tissue sections were permeabilized in a Labec mini bench top incubator at 37°C for 9 minutes prior to overnight reverse transcription at 42°C (20-24h). Tissue was removed from the surface of the STOmics chip with tissue release buffer for 10 minutes at 55°C. cDNA was released and collected overnight at 55°C (16-18h). Resultant cDNA was purified using 0.8X AMPure XP beads (Beckman Coulter, CA, USA) and eluted in 42μl of nuclease-free water (KIT). cDNA was amplified using specific primers with the protocol: 95°C for 5 minutes, 15 cycles of 98°C for 20s, 58°C for 20s, and 72°C for 3 minutes, followed by 72°C for 5 minutes and a 12°C hold. A final cleanup at 0.6X was performed with amplified cDNA eluted in 40μl of TE buffer. Purified cDNA was taken through for CRISPRclean or library preparation.

### STOmics-GenX

Genomic and housekeeping gene depletion was conducted using the CRISPRclean single cell boost kit (Jumpcode Genomics). Ribonucleoprotein (RNP) complex formation occurred by combining CRISPRclean 10X Cas9 buffer, RNase inhibitor, guide RNA, and Cas9 in a nuclease-free microcentrifuge tube followed by a 10-minute incubation at room temperature. A total of 50ng of purified cDNA was made up to 15μL with CRISPRclean nuclease-free water and added to the RNP alongside 1.5μL of CRISPRclean 10X Cas9 buffer. The resultant CRISPR digestion reaction was incubated at 37°C for 60 minutes followed immediately by a 2-minute incubation on ice. The CRISPR digestion reaction was made up to 50.3μL with CRISPRclean nuclease-free water and purified with AMPure beads at 0.6X. Purified CRISPRclean-depleted cDNA was eluted in 15μL of CRISPRclean nuclease-free water and taken through standard STOmics library preparation.

### Library preparation and sequencing

A total amount of 20ng of cDNA or 35ng of CRISPRclean-depleted cDNA was enzymatically fragmented at 55°C for 10 minutes. Once the reaction reached 4°C stop buffer was added, mixed thoroughly, and incubated at room temperature for 5 minutes. Barcodes were ligated to fragmented cDNA and amplified with the protocol: 95°C for five minutes, 13 cycles of 98°C for 20s, 58°C for 20s, and 72°C for 30s, followed by 72°C for five minutes and a 12°C hold. Samples were purified using a double size selection, first at 0.55X and then at 0.48X to obtain a target size range from 300-600bp. The final product was eluted in 20μl of TE buffer. To make DNBs for sequencing, libraries were denatured at 95°C for 3 minutes, 40°C for three minutes, then cooled to 4°C prior to the addition of Make DNB reaction mix 2. DNBs are synthesized in a thermal cycler over 30 minutes at 30°C. Stop DNB reaction buffer was added and carefully mixed with a wide bore tip on ice. Sequencing was then performed on the DNBSEQ-G400RS with both samples (STOmics and STOmics-GenX) sequenced on one flow cell using the DNBSEQ-G400RS FCL PE100 High-throughput Sequencing kit.

### Data processing

Raw sequencing reads were processed utilizing SAW v5.1.3 (BGI, https://github.com/BGIResearch/SAW). A STAR genome index was created using the Homo_sapien.GRCh38 primary assembly and Homo_sapien.GrCh38.106 annotation file (STAR v2.7.10a) and utilized for mapping by SAW. Reads were filtered to have a valid co-ordinate identity (CID) with a mismatch of 2. A count file was made by quantifying annotated reads after de-duplication based on CID, gene ID, and UMI. Multi-mapped read correction was included, as per SAW v5.1.3.

### Data analysis

Data was analysed at bin50 using Seurat v4.0.0 following SCTransform with variable.features.n = 2000 and return.only.var.genes = FALSE. Annotation was performed by summing the SCT counts of canonical genes: hepatocyte/epithelial - *ALB, KRT8, KRT18, KRT19, KRT5, KRT14*; myeloid - *LYZ, CD163, FOLR2, TREM2, KIT, SIGLEC, S100A8, S100A9, C1QA, CLEC10A*; fibroblast - *ACTA2, TAGLN, THY1, MYH11*; endothelial - *PECAM1, VWF, PLVAP, ACKR1*; and lymphoid - *CD3E, CD3D, CD8A, CD8B, CD4, IL7R, CXCL13, TRAV, TRBV, GNLY, NCAM1, MZB1, CD79A, CD19*. As the data at bin50 contains several cells, Fig. 2A assigned annotation from the minority to majority cell type (lymphoid, endothelial, fibroblast, myeloid, then hepatocyte/epithelial). Signature/gene expression feature plots for STOmics and STOmics-GenX utilize the same scale, with the max cutoff equal to the lowest maximum UMI.

## Data availability

The raw sequencing data of Stereo-seq is available at https://www.dropbox.com/sh/adri4lfdp7ikj6z/AADi-8S5RnzhVcoEHmjEJa4ta?dl=0. The codes are available at https://github.com/sharmaalab/STOmics-GenX

## Acknowledgements

This work is supported by start-up funds from the Harry Perkins Institute of Medical Research and Curtin University and an Ideas grant (2010795) from the National Health and Medical Research Council (NHMRC) awarded to A.S.

## Author contributions

A.S. and L.M. conceptualized the study and supervised the project; J.C. conducted the experiments and analysed data with contribution from R.P., S.G., C.K., Jo.C.; X.S., J.A., K.W., S.P., B.Y., J.P. contributed resources; L.Q. and J.G. collected clinical samples; J.C. generated figures; J.C. and A.S. wrote the manuscript, with all authors contributing to writing and providing feedback.

