## Supplementary Figure Legend for "STOmics-GenX: CRISPR based approach to improve cell identity specific gene detection from spatially resolved transcriptomics"

### Supplementary Figure Leagends:

**Fig S1. STOmics-GenX impacts highly variable genes. (A)** The residual variance is similar by methodology, yet an additional 592 HVGs were detected at a residual variance above 1.4 by STOmics-GenX. Of the HVGs, 719 HVGs had a residual variance above 1.4 in STOmics only **(B)**, with 1,311 only in STOmics-GenX **(C)**, leaving 1,015 present in both **(D)**. Mitochondrial and ribosomal genes are highlighted in red.
