## Supplementary figures and images for "STOmics-GenX: CRISPR based approach to improve cell identity specific gene detection from spatially resolved transcriptomics"

### Supplementary Figure

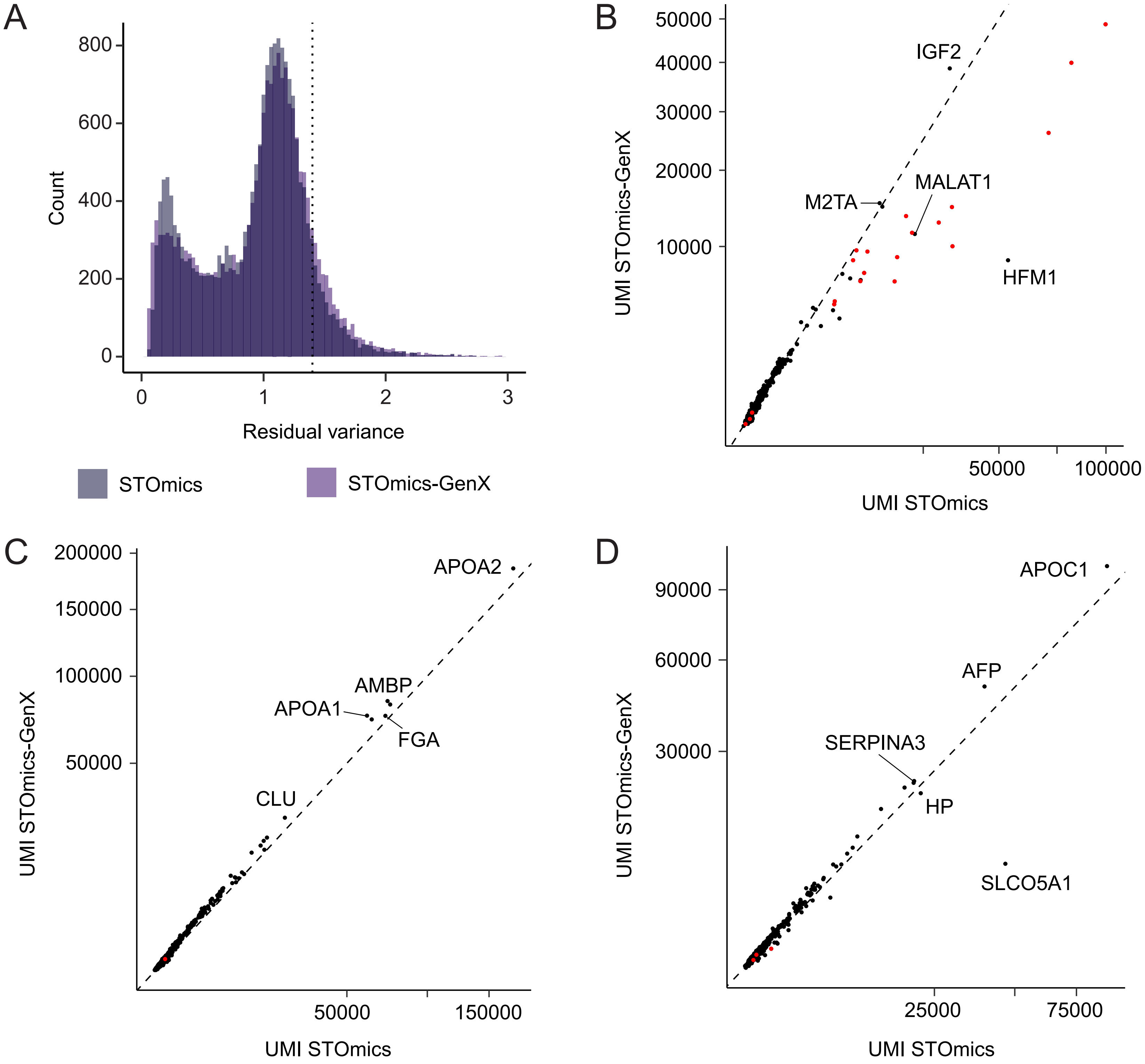
